# Enhancing Detection of Polygenic Adaptation: A Comparative Study of Machine Learning and Statistical Approaches Using Simulated Evolve-and-Resequence Data

**DOI:** 10.1101/2024.11.28.625827

**Authors:** Cosima Caliendo, Susanne Gerber, Markus Pfenninger

## Abstract

Detecting signals of polygenic adaptation remains a significant challenge in population genomics, as traditional methods often struggle to identify the associated subtle, multi-locus allele-frequency shifts. Here, we introduced and tested several novel approaches combining machine learning techniques with traditional statistical tests to detect polygenic adaptation patterns in time-series of allele frequency changes from whole genome data. We implemented a Naive Bayesian Classifier (NBC) and One-Class Support Vector Machines (OCSVM), and compared their performance against the classical Fisher’s Exact Test (FET). Furthermore, we combined machine learning and statistical models (OCSVM-FET and NBC-FET), resulting in 5 competing approaches. Using a simulated data set based on empirical evolve-and-resequencing *Chironomus riparius* genomic data, we evaluated methods across evolutionary scenarios, varying in generations, selection strength and numbers of loci under selection. Our results demonstrate that the combined OCSVM-FET approach consistently outperformed competing methods, achieving the lowest false positive rate, highest area under the curve, and high accuracy. The performance peak aligned with what we term the ‘late dynamic phase’ of adaptation—the period after initial selection has occurred but before fixation—highlighting the method’s sensitivity to ongoing selective processes and thus its value for experimental approaches. Furthermore, we emphasize the critical role of parameter tuning, balancing biological assumptions with methodological rigor. Our approach offers a powerful tool for detecting polygenic adaptation from time series, e.g. pool sequencing data from evolve-and-resequence experiments.

## Introduction

Rapid phenotypic adaptation is a key phenomenon in nature, allowing organisms to quickly adjust to changing environments [1]. The speed and extent of adaptive phenotypic changes can vary widely across taxa and environmental contexts, from subtle alterations in gene expression to dramatic morphological shifts [2]; for instance, early studies have documented rapid adaptive responses such as heritable wing size evolution in fly populations [3]. Beyond these discrete adaptive events, populations can also exhibit so-called adaptive tracking, where they continuously adjust traits through phenotypic plasticity or rapid microevolutionary shifts in response to fluctuating environmental conditions. Examples include beak morphology changes in Galápagos finches during drought period [4] and population-level responses deomstrated in both natural and experimental *Drosophila* and *Chironomus* populations [5] [6]. While such adaptive shifts of phenotypic traits can be more or less readily observed, detecting and attributing their underlying genomic signatures remains challenging [7]. This is particularly the case for polygenic traits where adaptive phenotypic changes are due to many loci with small individual effects [8]. This process plays a crucial role in species survival and diversification, particularly in the face of anthropogenic changes and climate fluctuations [9] [10].

Rapid adaptation can occur through selection on standing genetic variation and *de novo* mutations [11]. While some adaptive traits are controlled by single genes with large effects, many phenotypic changes in quantitative traits involve selection acting on multiple genes simultaneously, a process termed polygenic adaptation [12]. As polygenic adaptation does not necessarily require fixation of alleles or even large allele frequency changes it is often challenging to detect and measure the individual contributions of loci under selection [13]. Understanding the processes of rapid adaptation and its genomic basis is not only of fundamental scientific interest but has important societal implications for nature conservation biology, agriculture, climate change mitigation and medicine [14].

In recent years, the advent of high-throughput sequencing technologies has enabled researchers to generate vast amounts of genomic data, presenting both opportunities and challenges for detecting genomic signatures of adaptation [15]. These technologies have revolutionized our ability to study genetic variation at unprecedented scales, from single nucleotide polymorphisms (SNPs) to structural variants across entire genomes [16]. This wealth of data has led to the development of new statistical methods and bioinformatic tools for identifying genomic regions under selection [17]. However, the sheer volume and complexity of genomic data also present significant analytical and computational challenges, particularly in distinguishing true signals of molecular adaptation from background noise [18].

Pool sequencing (Pool-Seq) provides cost-effective population-level allele frequency estimates by sequencing pooled DNA from multiple individuals [19] [20] [21]. Successfully applied in contexts ranging from local adaptation studies in *Drosophila* [22] to experimental evolution, Pool-Seq enables broader temporal and spatial sampling than individual sequencing for equivalent resources. While both Pool-Seq and individual short-read approaches face analytical challenges - including haplotype phasing limitations and reduced power for rare variant detection [21] - Pool-Seq excels at the allele frequency estimation central to detecting polygenic adaptation [23].

While our understanding of rapid phenotypic adaptation has grown, the underlying molecular dynamics and genomic architectures driving these adaptations in many species remain an active area of research. Significant progress has been made in model organisms like *Drosophila* [24], *Chironomus* [25], *Arabidopsis* [26] und *Fagus* [27] providing valuable insights into the genetic basis of adaptation [24] [25]. These studies have revealed complex patterns of polygenic adaptation, where multiple loci contribute to adaptive traits [28].

Despite these advances, significant challenges remain. First, the broader application of findings from model organisms to diverse taxa remains difficult, as genetic architectures and selective pressures can vary substantially across species. Second, while some methods have been developed to detect both hard and soft selective sweeps in individual sequencing data [29] [30], traditional approaches for detecting selection have largely focused on identifying strong selective sweeps or large-effect loci [31], which represent only a probably minor subset of selection processes. Furthermore, these existing methods, including those for soft sweeps, are primarily designed for individual sequencing data and cannot be directly applied to Pool-Seq data. This limitation is particularly problematic for time series data, where subtle but coordinated allele frequency shifts across multiple loci may indicate important adaptive responses.

Existing approaches for detecting selection signals can be broadly categorized into three groups: (1) statistical tests on observed changes in allele frequency, (2) allele frequency spectrum-based summary statistics, and (3) machine learning methods. Each category offers certain advantages but also presents limitations when applied to polygenic adaptation scenarios.

Statistical tests of allele frequency changes, such as Fisher’s Exact Test (FET) and the Cochran–Mantel–Haenszel (CMH) test, are commonly employed to identify significantly altered allele frequencies between timepoints. These approaches are particularly suitable for evolve-and-resequence experiments using Pool-Seq data [32] [24] [33] [34] [3]. However, when used alone, these tests are known to produce p-value inflation—an excess of low p-values compared to the expected uniform distribution under the null hypothesis—leading to elevated false-positive rates [35]. Conversely, applying overly stringent significance thresholds can bias detection toward strong selective events, potentially missing the subtle signals characteristic of polygenic adaptation.. Conversely, applying overly stringent significance thresholds can bias detection toward strong selective events, potentially missing the subtle signals characteristic of polygenic adaptation.

Summary statistics based on nucleotide diversity patterns represent a second approach for detecting selection. Tools like PoPoolation2 [36] facilitate the calculation of metrics such as Tajima’s D, π, and Fst, offering flexibility in analysis scale from predefined windows to base-by-base calculations. However, these methods face challenges in determining the optimal scale for analysis: large windows may obscure subtle changes, while narrow windows might overemphasize statistical fluctuations [13]. Additionally, many summary statistics were developed primarily to detect classic selective sweeps rather than polygenic adaptation, where coordinated small shifts across multiple loci are the expected pattern. Furthermore, summary statistics are most powerful for detecting completed selection in the past [37], whereas detecting ongoing polygenic adaptation requires sensitivity to transient, directional frequency changes before equilibrium is reached. Therefore, there is a pressing need for novel data analysis approaches that can effectively detect and characterize polygenic selection, especially in longitudinally sampled genomic data.

In recent years, machine learning approaches have shown promise in addressing complex pattern recognition problems in genomics. Several studies have applied supervised learning methods to detect selective sweeps [38] [39]□HYPERLINK “https://doi.org/10.1093/molbev/msab332” [40], but these typically require labeled training data of known selected regions, which are rarely available for most non-model organisms, or require more data-specific information as for example haplotypes or recombination graphs. Furthermore, most existing machine learning applications in selection detection have focused primarily on classic selective sweeps rather than polygenic adaptation patterns., but these typically require labeled training data of known selected regions, which are rarely available for most non-model organisms, or require more data-specific information as for example haplotypes or recombination graphs. Furthermore, most existing machine learning applications in selection detection have focused primarily on classic selective sweeps rather than polygenic adaptation patterns.

To address these limitations, we propose and test novel combinations of machine learning techniques and statistical tests specifically designed to detect polygenic adaptation signals. Our approach leverages the pattern recognition capabilities of unsupervised learning methods—One-Class Support Vector Machines (OCSVM) [41] and Naive Bayesian Classifiers (NBC) [42], —combined with traditional statistical testing. Unlike supervised methods, these algorithms do not require pre-labeled examples of selection, making them particularly suitable for exploratory analysis of genomic data where ground truth is in general initially unknown.

OCSVM has proven effective at identifying collective, unusual patterns across diverse fields, from detecting anomalous heart rhythms in ECG data [43] to identifying environmental anomalies in atmospheric measurements [44]. As an unsupervised learning method, OCSVM’s ability to detect anomalous patterns without requiring pre-labelled training data makes it particularly promising for identifying adaptive loci in allele frequency changes, where the adaptive variants are, by definition, initially unknown.

NBC applies Bayes’ theorem to classify data based on the probability of features occurring together, assuming independence between predictors [45] [46]□HYPERLINK “https://doi.org/10.4018/978-1-59904-849-9.CH130”. In our context, assuming that most loci evolve in a neutral or nearly neutral fashion, NBC learns the probability distribution of summary statistics under neutral evolution and calculates the likelihood that observed genomic patterns belong to this neutral class. NBC provides an additional advantage by offering a probabilistic framework for classification, aligning well with the stochastic nature of evolutionary processes and allowing for nuanced interpretation of results.. NBC provides an additional advantage by offering a probabilistic framework for classification, aligning well with the stochastic nature of evolutionary processes and allowing for nuanced interpretation of results.

Both OCSVM and NBC demonstrate flexibility in pattern recognition, capable of capturing complex, non-linear relationships in the data, making them well-suited to detect subtle, polygenic adaptation patterns that might be missed by linear statistical methods. These classifiers are also suited for handling high-dimensional data, a common characteristic of genomic datasets, potentially uncovering patterns that are not apparent in univariate analyses. Contrary to window-based methods, these classifiers can analyse data across the genome without relying on arbitrary window sizes, potentially capturing both localized and broad-scale patterns of adaptation. By integrating machine learning with statistical testing in our OCSVM-FET and NBC-FET approaches, we aim to leverage the strengths of both paradigms: the pattern recognition capabilities of machine learning algorithms and the statistical rigor of traditional methods. We evaluate these novel approaches against conventional FET using simulated data with known selection patterns, providing a controlled environment to assess performance across various evolutionary scenarios and selection intensities.

## Material and Methods

### Simulations

To simulate polygenic adaptation, we employed MimicrEE2 (version 2.06), a forward-in-time simulator operating in quantitative trait (QT) mode, which models selection on a quantitative trait influenced by multiple loci. We generated haplotype input files from 66 *C. riparius* individuals sequenced across 712 scaffolds (8,284,035 SNPs total), which served as the ancestral generation (G0) for our simulations.

We designed a full factorial experimental setup exploring three key variables: (1) time under selection (10, 20, 40, and 60 generations), (2) number of loci under selection (10, 50, 100, 250, and 500 loci), and (3) selection strength (weak, medium, strong). For each combination of these factors, we ran 10 independent replicates, resulting in 600 total simulation scenarios (4 × 5 × 3 × 10).

#### Population Parameters

The simulated population consisted of 1,000 diploid individuals per generation following a discrete-generation Wright-Fisher model. Selection acted on a quantitative trait with broad-sense heritability (H²) of 0.8, meaning 80% of phenotypic variance was attributable to additive genetic effects and 20% to environmental noise.

#### Selection Regime

We implemented truncating selection on the phenotypic trait, where individuals with phenotypes below a fitness threshold were excluded from reproduction. Selection strength varied across three regimes:

- **Weak selection:** Truncation at the 20th percentile (bottom 20% excluded), corresponding to a shift in the phenotypic optimum of approximately 0.35 Haldanes (phenotypic standard deviations) from the ancestral population mean
- **Medium selection:** Truncation at the 50th percentile (bottom 50% excluded), corresponding to a shift of approximately 0.80 Haldanes
- **Strong selection:** Truncation at the 80th percentile (bottom 80% excluded), corresponding to a shift of approximately 1.40 Haldanes

This design created a gradient of selection intensities while maintaining constant directional selection toward higher phenotypic values.

#### Genetic Architecture

Loci under selection were randomly selected from the genome, requiring a minimum allele frequency of 0.15 in the ancestral population to ensure sufficient standing genetic variation. Effect sizes for selected loci were drawn from a uniform random distribution ranging from 1.5 to 3.5 phenotypic units, creating heterogeneous genetic architectures where loci contributed variably to the trait.

Phenotypic values for each individual were calculated as:

**P = G + E**

where G represents the genetic component (sum of allelic effects across all selected loci, determined by genotype and effect sizes) and E represents environmental noise sampled from a normal distribution with variance scaled to achieve the specified heritability of 0.8.

#### Recombination

Recombination rates were modeled across the 712 scaffolds, with rates for each scaffold drawn uniformly at random between 0.1 and 4.0 cM/Mb, reflecting natural variation in recombination across the *C. riparius* genome [48].

#### Simulation Design Rationale

This simulation design allowed us to model rapid adaptation scenarios with varying degrees of genetic architecture complexity, from oligogenic (10 loci) to highly polygenic (500 loci) adaptation, across different time scales and selection intensities. All simulated individuals from each generation were retained for downstream analysis without subsampling.

## Methods for detecting polygenic adaptation

Allele frequencies were calculated for each SNP position across all sampled generations using the R package poolSeq [Taus ref]. Throughout all analyses, allele frequencies were consistently calculated with respect to the reference allele as defined in the *C. riparius* genome assembly (Crip3.0), rather than polarizing by the derived or increasing allele. This means we did not fold the allele frequency spectrum, and frequency changes (Δp) represent changes in reference allele frequency, which can be either positive (reference allele increasing) or negative (reference allele decreasing) under selection.

### Fisher’s Exact Test (FET)

Fisher’s exact tests were conducted to compare allele frequencies between the ancestral (first) generation and generation 10, 20, 40, and 60, yielding corresponding p-values using poolSeq (version 1.15.4, [49]) with R (version 4.3.2) [49]. Results were corrected for multiple comparisons using the Benjamini-Hochberg correction using the stats R package (version 4.5.0). We employed a stringent significance threshold (adjusted p-value < 0.001) as proposed by Pfenninger et al. [50] for detecting strong selection signals while minimizing false positives. This conservative cutoff was chosen deliberately to identify loci with substantial allele frequency changes that are unlikely to occur through drift alone. While less stringent thresholds (e.g., 0.01 or 0.05) would identify more candidate loci, they would also increase the false discovery rate. For our primary goal of identifying robust polygenic adaptation patterns, prioritizing specificity over sensitivity was deemed appropriate.

### The One Class Support Vector Machines (OCSVM)

The OCSVM algorithm [41] was implemented using Python (version 3.8.13) and scikit-learn (version 1.1.3) [51] to identify potential signatures of polygenic adaptation in allele frequency data. Similar to our FET approach, we applied OCSVM to compare allele frequencies between the ancestral generation and each subsequent generation (10, 20, 40, and 60), ensuring methodological consistency across approaches.

OCSVM is specifically designed to detect anomalies or outliers [41]. in unlabeled data. The algorithm learns to identify the boundary that encapsulates the “normal” distribution of data points, classifying samples outside this boundary as anomalies. In our context, we consider allele frequency changes (AFCs) between generations as our data points, with the majority representing neutral genetic drift (the “normal” class), while outlier points represent loci potentially under selection.

The outlier criterion in OCSVM is determined by the decision function’s output: data points receiving negative scores (−1) are classified as outliers (potential selection candidates), while those with positive scores (+1) are classified as normal (likely neutral). This classification is based on the data point’s position relative to the decision boundary learned during training. Points falling outside the boundary (i.e., in low-density regions of the feature space defined by the summary statistics) are flagged as outliers, while those inside the boundary (in high-density regions) are classified as normal. The algorithm constructs this boundary in a high-dimensional space using the “kernel trick” to capture non-linear relationships in the data (Figure 1). We employed the Radial Basis Function (RBF) kernel, which is mathematically expressed as:

**Fig 1.**
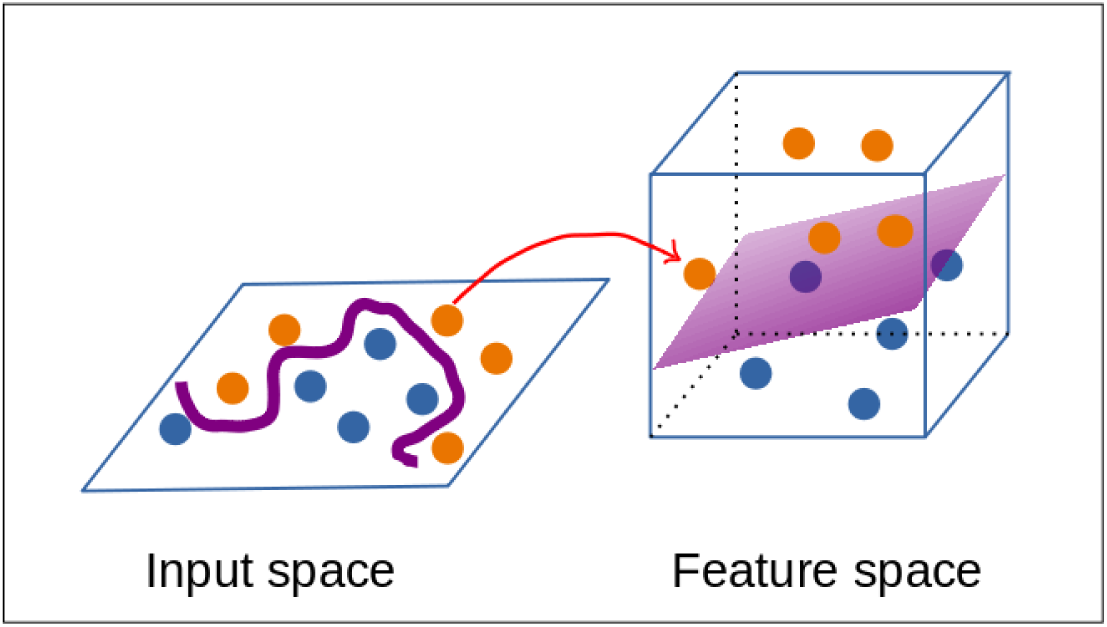
Schematic illustration of One Class Support Vector Machine’s *kernel trick.* The so-called kernel trick is used by the OCSVM algorithm to compute dot products between pairs of data points in higher-dimensional space

The RBF kernel is mathematically expressed as:

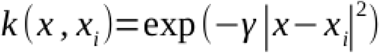

Two crucial parameters influence the algorithm’s behavior:

- γ (gamma): Determines the influence radius of each data point on the decision boundary. Lower values create a smoother, more generalized boundary, while higher values create a more complex boundary that can capture localized patterns.
- ν (nu)Controls the upper bound on the fraction of training errors that fall outside the learned boundary and the lower bound on the fraction of support vectors. Essentially, this parameter balances between maximizing the margin and minimizing misclassification of the training data [52].

These parameters were optimized through 5-fold cross-validation using previously published data on *C. riparius* [53]. We tested γ values ranging from 0.01 to 0.5 and ν values ranging from 0.01 to 0.2, ultimately selecting the optimal combination of ν = 0.01 and γ = 0.05 based on highest accuracy FPR and AUC (see section Parameter Fine Tuning). This optimization process ensured the model was calibrated specifically for detecting genetic selection signals in this species while minimizing false positives.

For each comparison between the ancestral and a subsequent generation, allele frequencies were extracted using the poolSeq R package and provided as input to the OCSVM algorithm. The output classification (−1 for outliers, +1 for normal) was used to identify candidate loci potentially involved in the adaptive response. By applying this approach consistently across all generational comparisons (generation 0 versus 10, 20, 40, and 60), we could track the emergence and persistence of selection signals throughout the evolutionary trajectory of the population.

### Naive Bayesian Classifier (NBC)

We employed a NBC algorithm [42] to distinguish between normal (non-anomalous) and anomalous data points based on their allele frequency changes in two-dimensional (2D) space. The NBC is a probabilistic classifier based on Bayes’ theorem, where the fundamental idea involves modelling the probability density functions of data set distributions, in this case for non-anomalous and anomalous data. Each data point is represented by two features: the initial allele frequency (generation 0) and the allele frequency at a later generation. Anomalous data represents candidate SNPs showing high allele frequency changes (AFC) between ancestral and adapted generations, either increasing (beneficial) or decreasing (disadvantageous), while non-anomalous data represents positions not targeted by selection. The classifier models these patterns using three probability distributions: one for neutral variation and two for selected variants. The mathematical function modelling these areas of interest is described in the supplement and was implemented using Python (version 3.8.13) and the library scipy (version 3.8.13) and the library scipy (version 1.7.3) [54].

The NBC implementation uses mean vectors (μ) and covariance matrices (Σ) to model three distinct classes in the data (see Supplement for details): two classes for anomalous patterns (allele frequencies changing from low to high or vice versa) and one for non-anomalous background patterns showing minimal changes. These parameters were optimized using published *C. riparius* data [50] as described above, to ensure reliable detection of genetic changes in this species.

### Combination with Fisher’s Exact Test

We developed a combined approach that integrates the results from 3 independent analyses: the binary classification results from OCSVM and NBC, along with the statistical hypothesis testing of FET. Using Python with the library *pandas*, we merged the previously obtained results from all three methods by matching genomic positions. This integration process identified overlapping positions – those flagged as anomalous by either the OCSVM and NBC approach, respectively, and simultaneously showing statistical significance in FET. This combined approach leverages the strengths of both machine learning and traditional statistical testing to identify and narrow down candidate positions targeted by polygenic adaptation.

### Parameter Fine Tuning

We optimized key parameters for both NBC (μ and Σ) and OCSVM (ν and γ) using a two-stage approach: initial tuning on empirical data followed by extensive validation on simulated datasets.

#### Initial Parameter Estimation

For initial tuning, we utilized genome-wide allele frequency data from Pfenninger et al. [53], which identified 10 independent loci showing signatures of selection in *C. riparius* during rapid adaptation. These 10 loci served as positive examples of selection, while we randomly sampled 9,000 SNP not flagged as selected from the entire genome to represent the non-selected class. This approach created a realistically imbalanced training scenario approximating the typically rare nature of selective events in genomes (see Supplementary Materials for detailed optimization procedures).

#### Cross-Validation Procedure

To address this substantial imbalance in our training data and ensure robust parameter estimation, we implemented a 5-fold cross-validation procedure. The dataset was split into 5 equal subsets, with each subset maintaining the original ratio of selected to neutral loci. For each fold, we trained the algorithms on 80% of the data and evaluated performance on the remaining 20%. This process was repeated five times, with each subset serving once as the validation set.

#### Parameter Grid Search

For OCSVM, we performed a grid search over ν values ranging from 0.001 to 1 (0.001, 0.005, 0.01, 0.013, 0.02, 0.1, 1) and γ values from 0.01 to 1 (0.01, 0.05, 0.5, 1). For NBC, we explored 10 different combinations of μ and Σ values, ensuring the resulting covariance matrices remained valid positive semi-definite (see Supplementary Fig S2 for detailed parameter combinations).

#### Performance Evaluation

For each parameter combination, we calculated multiple performance metrics: accuracy, false positive rate (FPR), true positive rate (TPR), and area under the ROC curve (AUC). Given the imbalanced nature of our data, we placed particular emphasis on the FPR and AUC metrics, as accuracy alone can be misleading when data is imbalanced because classifiers tend to be biased towards the majority class, leading to poor performance on the minority class [55] [56].

### Performance Metrics

We evaluated the five approaches (FET, OCSVM, NBC, OCSVM-FET, NBC-FET) using three complementary metrics implemented with Python (version 3.8.13) and scikit-learn: false positive rate (FPR), area under the receiver operating characteristic curve (AUC), and accuracy.

This combination of metrics was selected to provide a comprehensive performance assessment given our imbalanced dataset, where selected loci represent only a small fraction of all variants. While accuracy (proportion of correct classifications) offers an intuitive measure of overall performance, it can be misleadingly high in imbalanced datasets if a method primarily predicts the majority class. In contrast, AUC measures discriminative power by accounting for both sensitivity and specificity across different thresholds, making it less affected by class imbalance. FPR (ratio of false positives to all negative cases) specifically quantifies the tendency to misclassify neutral loci as under selection. Together, these metrics enabled robust comparison across detection methods and evolutionary scenarios [57]. Visualization was performed using matplotlib (version 3.5.2) [58]..

## Results

### Parameter Optimization on template data

Our parameter optimization process aimed to identify the optimal configuration for detecting subtle signals of polygenic adaptation while minimizing false positives. All performance metrics reported represent the mean values from 5-fold cross-validation on test data not used during training.

For the OCSVM algorithm with a radial kernel, we optimized two key parameters: ν, which influences the algorithm’s sensitivity to outliers, and γ, which controls the decision boundary curvature. Figure 2a illustrates the performance metrics across various parameter combinations. The optimal OCSVM configuration (ν=0.01, γ=0.05) achieved a balance between a low FPR (0.084), high AUC (0.937), and high accuracy (0.965).

**Fig 2.**
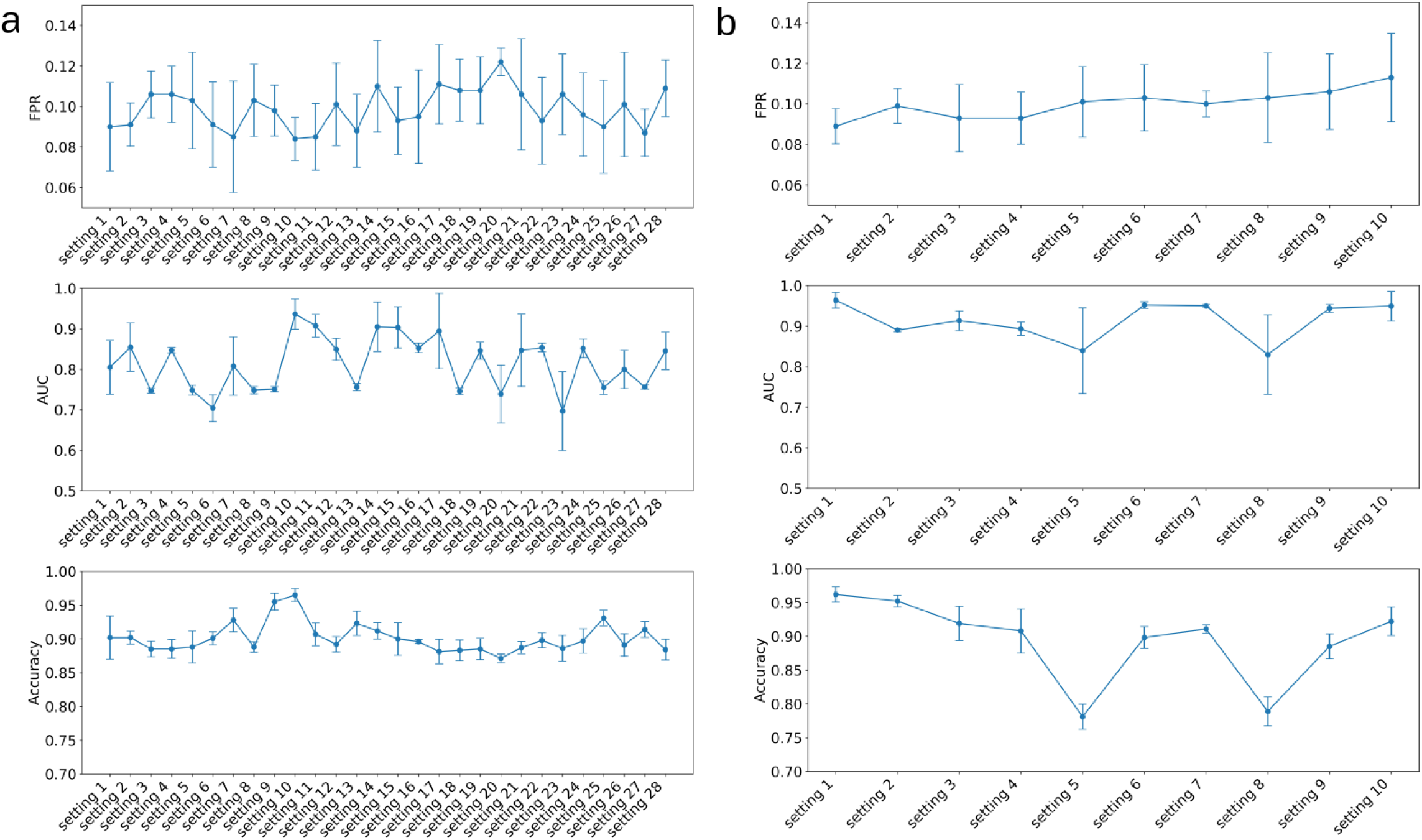
Parameter optimization for One Class Support Vector Machines (OCSVM) and Naive Bayesian Classifier (NBC) algorithms. (a) Performance metrics (False Positive Rate, Area Under the Curve, and Accuracy) for various parameter settings of the OCSVM with radial kernel. (b) Corresponding metrics for different NBC parameter configurations.

For the NBC, we optimized parameters defining three probabilistic models: one for the majority, non-anomalous data (represented by μ_1_ and Σ_1_) and two for the anomalous data (represented by μ_2_/μ_3_ and Σ_2_/Σ_3_). Figure 2b displays the performance metrics for 10 different parameter configurations. Setting 1 (μ_1_=[0.0, 0.0], Σ_1_=[0.15, 0.1], μ_2_/μ_3_=[0.1, 0.9], Σ_2_/Σ_3_=[0.25, 0.2]) provided the optimal performance with an FPR of 0.089, AUC of 0.964, and accuracy of 0.962.

While both methods achieved high performance metrics, we observed that NBC exhibited greater sensitivity to parameter changes, with performance varying substantially across different configurations (Table 1). In contrast, OCSVM showed more consistent performance across a wider range of parameter settings, suggesting higher robustness to parameter variations (for detailed information see Supplementary Tab S1).

**Tab 1.** Top 5 parameters settings for One Class Support Vector Machines (OCSVM) and Naive Bayesian Classifier (NBC)

| Setting | $\nu$ | $\gamma$ | FPR | AUC | ACCURACY |
| --- | --- | --- | --- | --- | --- |
| 10 | 0.01 | 0.05 | 0.084 | 0.937 | 0.965 |
| 11 | 0.01 | 0.5 | 0.085 | 0.908 | 0.907 |
| 14 | 0.013 | 0.05 | 0.11 | 0.905 | 0.912 |
| 15 | 0.013 | 0.5 | 0.093 | 0.903 | 0.9 |
| 17 | 0.2 | 0.01 | 0.111 | 0.895 | 0.881 |

b
| Seeting | $\mu_1$ | $\Sigma_1$ | $\mu_2/\mu_3$ | $\Sigma_2/\Sigma_3$ | FPR | AUC | ACCURAY |
| --- | --- | --- | --- | --- | --- | --- | --- |
| 1 | 0.0, 0.0 | 0.15, 0.1 | 0.1, 0.9 | 0.25, 0.2 | 0.089 | 0.964 | 0.962 |
| 10 | 0.0, 0.0 | 0.15, 0.1 | 0.7, 0.2 | 0.25, 0.2 | 0.113 | 0.945 | 0.922 |
| 6 | 0.0, 0.0 | 0.15, 0.2 | 0.3, 0.8 | 0.15, 0.1 | 0.103 | 0.952 | 0.898 |
| 7 | 0.0, 0.0 | 0.2, 0.15 | 0.35, 0.6 | 0.3, 0.2 | 0.1 | 0.95 | 0.91 |
| 3 | 0.0, 0.0 | 0.2, 0.1 | 0.3, 0.5 | 0.4, 0.2 | 0.093 | 0.913 | 0.991 |

(a) Top 5 parameter settings ( $\nu$ and $\gamma$ ) for the OCSVM algorithm with their respective performance metrics. (b) Top 5 parameter settings for NBC, showing $\mu$ and $\Sigma$ values for three approximations along with their performance metrics.

To assess generalizability beyond our empirical training data, we applied the optimized parameters to simulated data representing diverse evolutionary scenarios. Performance across these simulations

### Application on simulated data

To evaluate the performance of competing methods to detect patterns of polygenic adaptation where the ground truth is known, we utilized simulated data generated with Mimicree2 based on real *C. riparius* allele frequencies.

Our analysis of different approaches (FET, OCSVM, OCSVM-FET, NBC, and NBC-FET) across various generations (10, 20, 40, and 60) revealed that generation 40 provides the optimal balance for detecting loci under selection (Figure 3).

**Fig 3.**
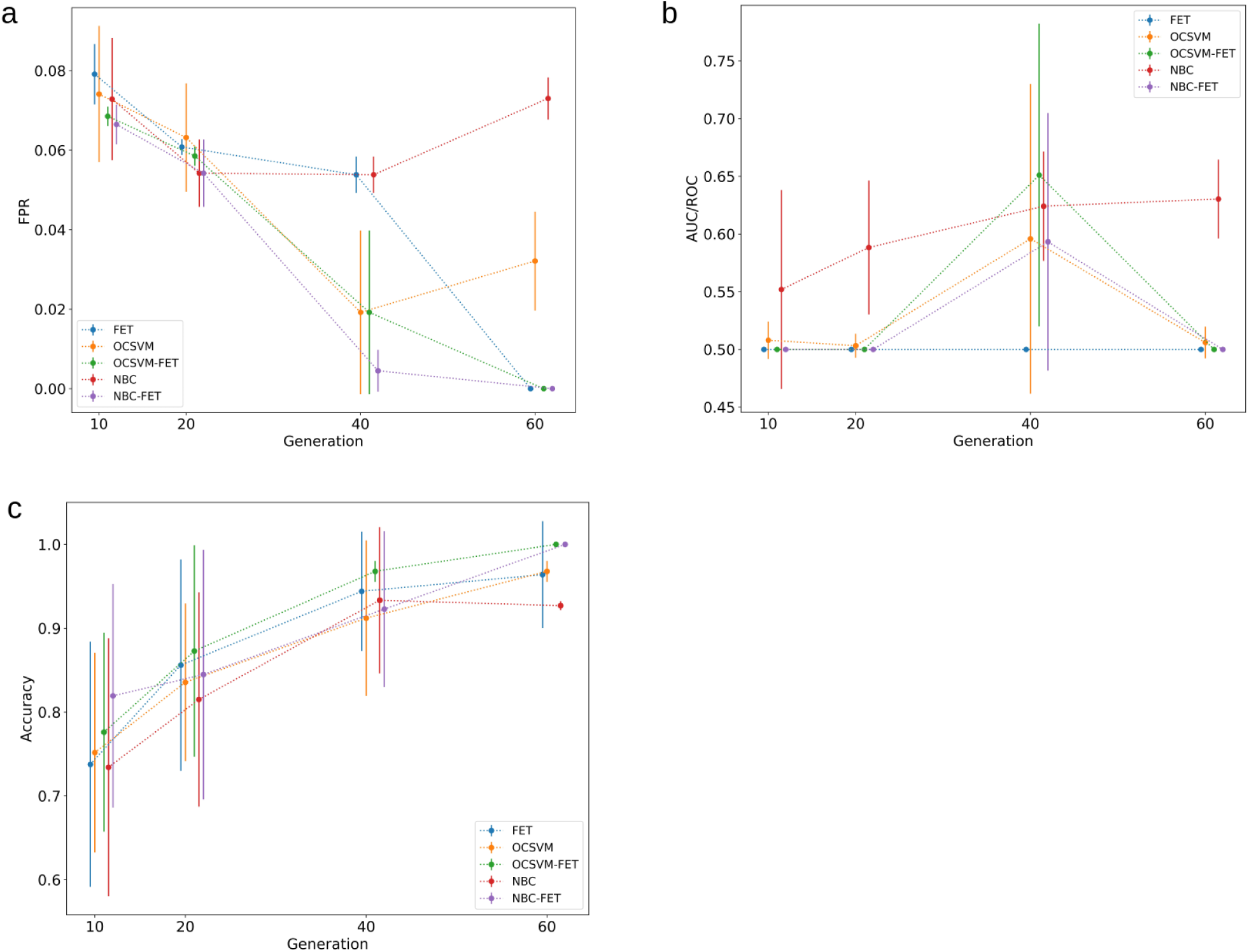
Performance comparison of five approaches for detecting loci under selection across different generations. The approaches evaluated are Fisher’s Exact Test (FET), One Class Support Vector Machines (OCSVM), Naive Bayesian Classifier (NBC) and the combination OCSVM-FET, NBC-FET. Performance metrics shown are (a) false positive rates (FPR), (b) area under curve/receiver operating characteristic (AUC/ROC), and (c) Accuracy. Data points represent mean values across simulations, with error bars indicating standard deviation. Generations tested: 10, 20, 40, and 60.

FPR showed a downward trend from generation 10 to generation 40 for all approaches (Figure 3a). NBC and OCSVM, showed a slight increase in generation 60, whereas the remaining approaches reached their lowest values in generation 60. AUC/ROC (Figure 3b) indicated that all approaches, except FET, reached peak performance at generation 40. Notably, NBC maintained fairly good performance (AUC>60%) even at generation 60, while other approaches declined to approximately 50%. FET consistently showed values around 50%, indicating random performance, while OCSVM-FET demonstrated the best overall results, particularly at generation 40. Accuracy (Figure 3c) showed a consistent improvement for all approaches with increasing generations, reaching a plateau-like signature from generation 40 to generation 60. OCSVM-FET achieved the highest accuracy at generation 40, followed by FET and NBC.

In summary, for the tested evolutionary scenarios, generation 40 emerged as the optimal scenario for all approaches, considering both accuracy and AUC. While 3 (OCSVM, OCSVM-FET and NBC-FET) of the 5 approaches reached their lowest values of FPR in generation 60, generation 40 shows comparable low values for these 3 approaches and additionally the lowest values for the remaining 2 approaches (FET, NBC).

To understand these results, we examined the allele frequency distributions across different generations (Figure 4), as well as phenotypic values. (Figure 5). Allele frequencies of SNPs identified to be under selection (red dots) became increasingly distinct from the background (blue dots) as generations progressed, with a clear separation emerging around generation 40 (Figure 4a). This pattern closely resembled the distribution observed in the real-life data used for parameter tuning (Figure 4b). While late-generation simulations (Fig. 4a, generations 40-60) show more pronounced fixation than typically observed in nature, our method excels at detecting selection signals in intermediate generations before complete fixation, which more accurately reflects real evolutionary processes.

**Fig 4.**
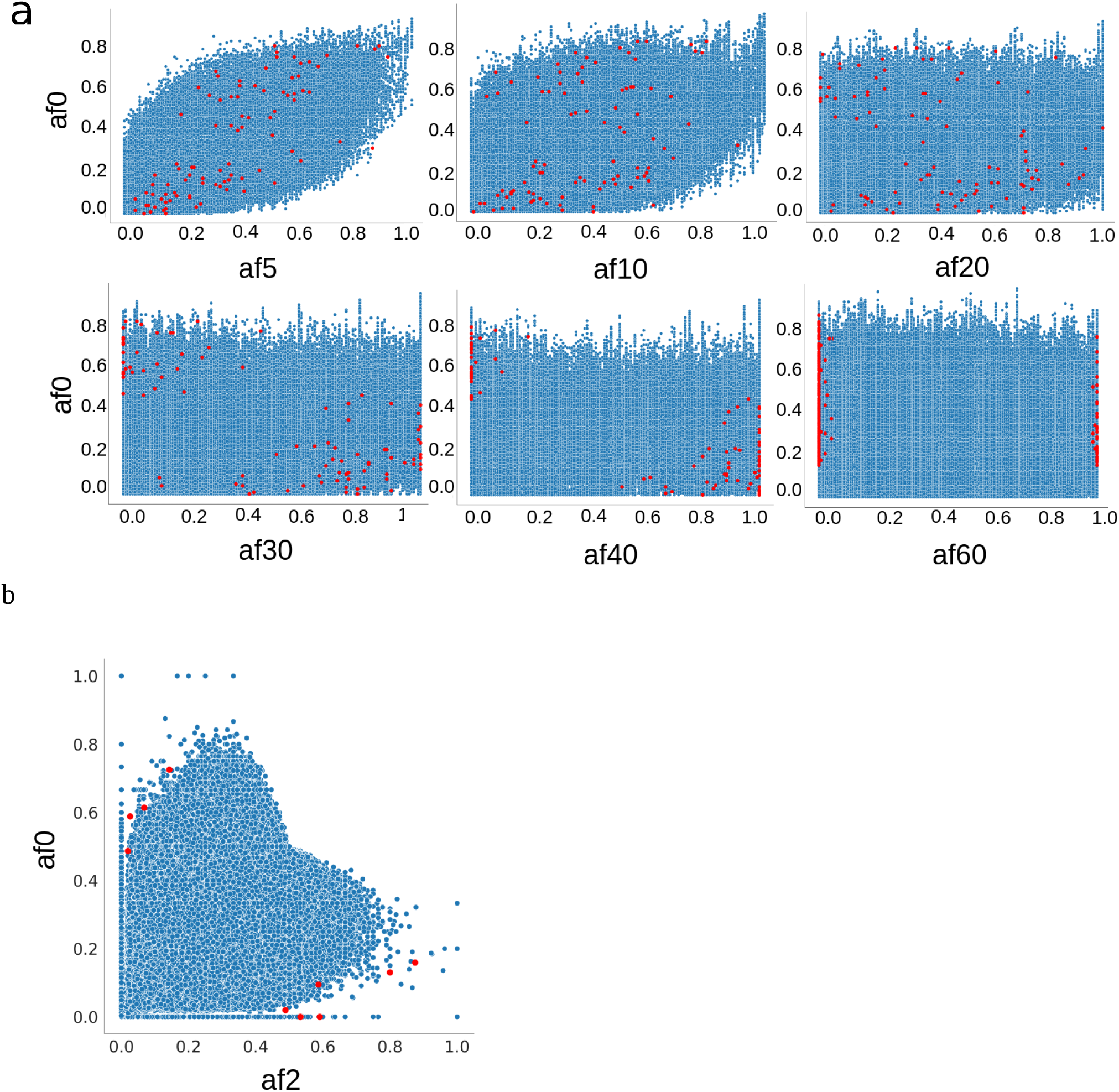
Allele frequency distributions across generations. (a) Simulated data showing allele frequencies (AF) at generations 5, 10, 20, 30, 40, and 60 plotted against initial allele frequencies (af0). Red dots represent selected SNPs, while blue dots represent background SNPs. (b) Real-life data used for parameter tuning, showing allele frequencies at generation 2 plotted against initial frequencies (af0). The distribution of anomalies identified loci in generation 40 of the simulated data most closely resembles the real-life data.

**Fig 5.**
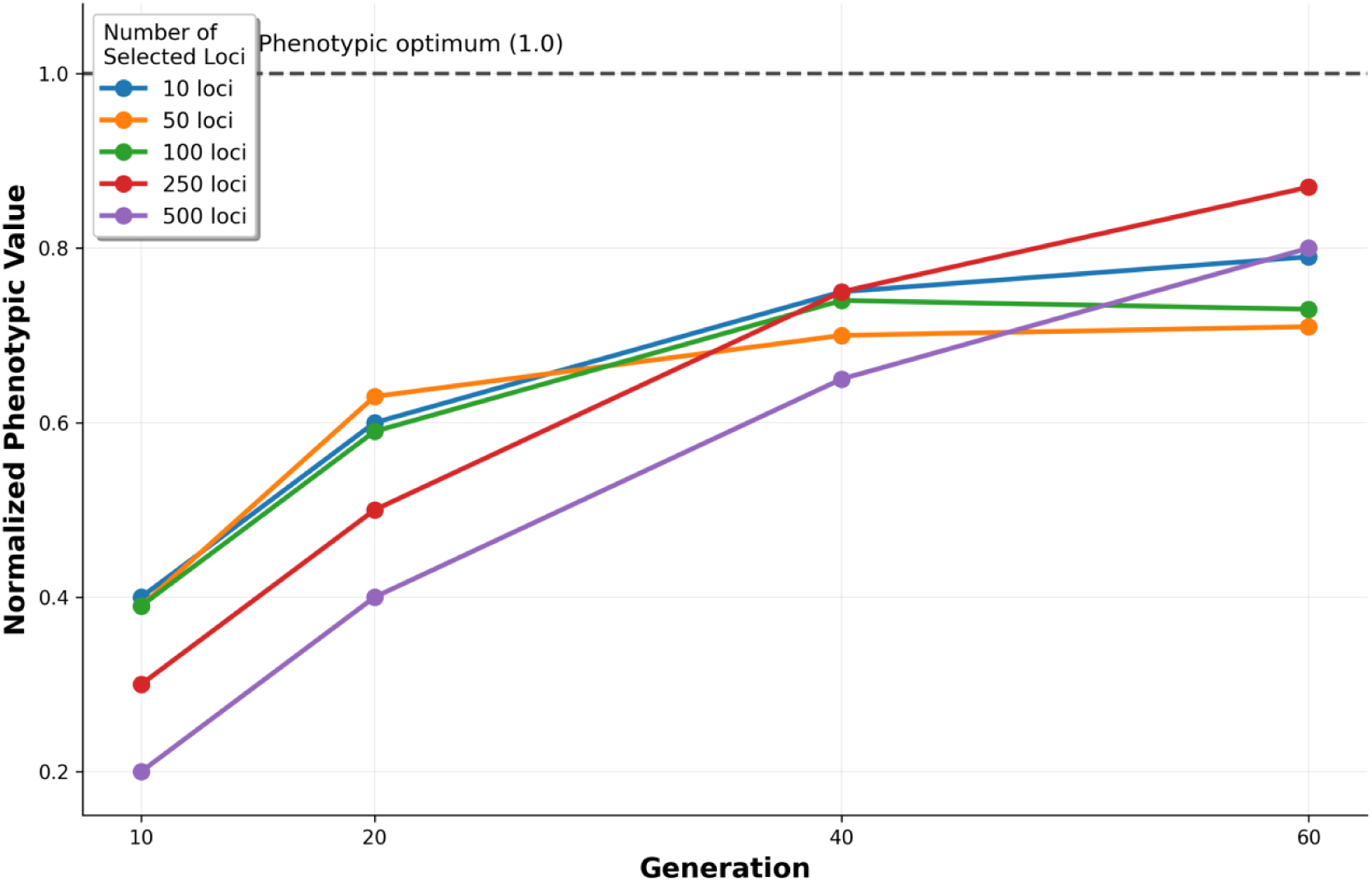
Normalized phenotypic trajectories over generations for different numbers of selected loci under strong selection. Values represent normalized phenotypic scores scaled relative to each scenario’s ancestral population mean (0) and theoretical evolutionary optimum (1.0, representing complete fixation of all beneficial alleles). Each point shows the mean normalized phenotypic value across 10 independent simulation replicates for populations with 10, 50, 100, 250, or 500 loci under selection. The dashed horizontal line at 1.0 indicates the normalized phenotypic optimum. Measurements were taken at generations 10, 20, 40, and 60. Note that while the normalization enables comparison of relative adaptive progress across scenarios, absolute phenotypic values differ among scenarios due to variation in the number of loci and their associated effect sizes. Simulations were conducted under strong truncating selection (80th percentile threshold, corresponding to a 1.40 Haldane shift) with heritability of 0.8.

The normalized phenotypic trajectories for different numbers of selected loci revealed distinct patterns of adaptation over 60 generations under strong selection (Fig. 5). Simulations with fewer loci (10 and 50) showed rapid initial increases in normalized phenotypic values, reaching approximately 0.60-0.63 by generation 20 but subsequently plateauing or even declining slightly (10 loci: 0.79 at generation 60; 50 loci: 0.71 at generation 60). This plateau effect suggests rapid depletion of standing genetic variation as beneficial alleles at the limited number of selected loci quickly approach or reach fixation.

In contrast, simulations with larger numbers of loci (250 and 500) displayed slower initial increases but continued to rise steadily throughout the 60-generation period. The 250-loci scenario achieved the highest normalized phenotypic value by generation 60 (0.87), approaching but not reaching the theoretical optimum (1.0). The 500-loci scenario showed the slowest initial response, with a normalized value of only 0.20 at generation 10, but exhibited sustained adaptive progress throughout the simulation period, reaching 0.80 by generation 60.

Notably, none of the simulated scenarios achieved the normalized optimum value of 1.0 within 60 generations, indicating that complete fixation of all beneficial alleles requires extended evolutionary time, even under strong directional selection. The intermediate architecture (250 loci) showed the best balance between rapid initial response and sustained long-term progress, suggesting an optimal range of genetic complexity for efficient polygenic adaptation under the simulated conditions.

Our OCSVM-FET approach detects polygenic adaptation by identifying collective patterns in allele frequency changes across multiple loci. Unlike methods that analyze each SNP independently, OCSVM captures genome-wide distribution patterns, recognizing coordinated shifts that may be subtle at individual loci but significant in aggregate. This enables detection of diffuse selection signals across many loci without requiring complete fixation at any single position. The method performs optimally with intermediate loci numbers (100-250), capturing polygenic architectures while maintaining statistical power to distinguish selection from background variation.

### OCSVM-FET shows Superior Performance Across Loci Numbers

Using generation 40 as the optimal timepoint, we evaluated the performance of different approaches across varying numbers of loci under selection (n = 10, 50, 100, 250, and 500) (Figure 6). FPR analysis showed consistent results of approximately 0.06 for FET and NBC across all loci settings. NBC-FET maintained a low FPR of around 0.01, while OCSVM and OCSVM-FET showed slightly elevated values for n=100, but otherwise maintained values around 0.02.

**Fig 6.**
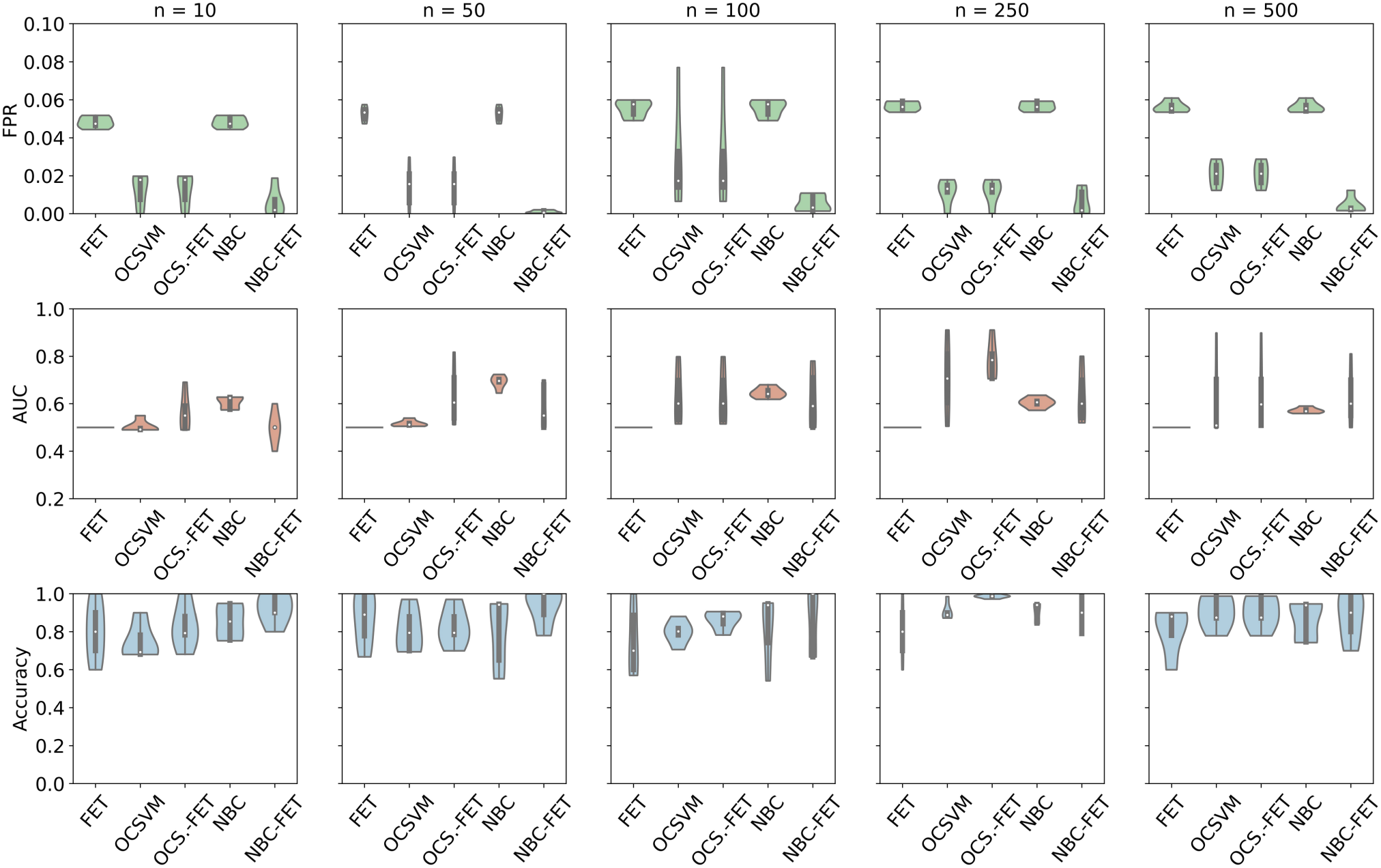
Comparative analysis of detection approaches across varying numbers of loci under selection at generation 40. The graph displays false positive rate (FPR) (first column graphs, green), area under curve/receiver operating characteristic (AUC/ROC) (second column graphs, red), and accuracy (third column graphs, blue for Fisher’s Exact Test (FET), One Class Support Vector Machines (OCSVM), Naive Bayesian Classifier (NBC) and the combined approaches OCSVM-FET (OCS.-FET), NBC-FET. Each row represents the number of loci under selection (n = 10, 50, 100, 250, 500).

AUC performance varied among approaches. FET consistently showed values around 50% across all loci settings, indicating random performance. OCSVM and OCSVM-FET demonstrated increasing performance with higher numbers of selected loci, peaking at n=250 (OCSVM >70% AUC, OCSVM-FET >80% AUC), followed by a slight decrease at n=500 (OCSVM <65% AUC, OCSVM-FET <70% AUC). NBC reached its optimal performance at n=50 (∼65% AUC), gradually declining to around 60% with increasing number of loci. Conversely, NBC-FET showed improving performance with increasing n, plateauing at n=100 with values consistently above 60% (Figure 6).

Accuracy analysis revealed that the OCSVM-FET approach achieved optimal results with n=250 selected loci, reaching approximately 99% accuracy. Accuracy generally increased with n for this approach, with a slight decrease to 80% at n=500. NBC-FET maintained high accuracy (around 90%) for n=10, 50, 100, and 150, with a minor decrease at n=500. FET showed no distinct trend, maintaining around 80% accuracy across various n values. OCSVM and NBC exhibited a gradual increase in accuracy with increasing n, both peaking at n=250 before slightly decreasing.

While focusing on generation 40, we also examined the performance of our methods across generations 10, 20, and 60 to provide a comprehensive view of how the detection of polygenic adaptation signals evolves over time (Supplementary Figs S3-S5).

At the early stage of generation 10 (Fig S3), all methods showed relatively poor performance across all loci numbers. The AUC/ROC values were generally lower compared to later generations, with most methods hovering around 50-60%, indicating only slightly better than random performance. Accuracy was also generally lower, with high variability across methods and loci numbers. For generation 20 (Fig S4), performance improved for most methods compared to generation 10, particularly for scenarios with higher numbers of loci under selection (n=250, 500). OCSVM and OCSVM-FET began to show better AUC/ROC values, especially for n=250 and n=500, though not as high as observed at generation 40. Accuracy also improved, with OCSVM-FET and NBC-FET showing the most consistent performance across loci numbers. Interestingly, for generation 60 (Fig S5), performance for most methods slightly declined or plateaued compared to generation 40, particularly for scenarios with higher number of loci (n=250, 500). This trend was most noticeable in the AUC/ROC and accuracy measures for OCSVM and OCSVM-FET. FPR remained relatively stable across generations for most methods. This decline in detection performance at later generations reflects weakening directional selection as populations approach the phenotypic optimum. Near the optimum, genetic redundancy allows compensatory allele frequency changes across loci to maintain the optimal phenotype, creating frequency fluctuations that increasingly resemble neutral drift despite ongoing stabilizing selection. This reduces the discriminability of selected loci from the genomic background, a characteristic feature of polygenic traits at their adaptive optimum.

### Optimal Framework Performance

Our comprehensive analysis indicated that the OCSVM-FET approach performed optimally when populations were intermediate in their adaptive trajectory toward the phenotypic optimum. In our simulations under strong selection (1.40 Haldane shift), this corresponded to generation 40 with intermediate polygenic architectures (250 selected loci) (Figure 7). Under these conditions, OCSVM-FET demonstrated consistently strong performance metrics: it achieved the lowest False Positive Rate, produced AUC values exceeding 80%, and maintained near-perfect accuracy of approximately 99%.

**Fig 7.**
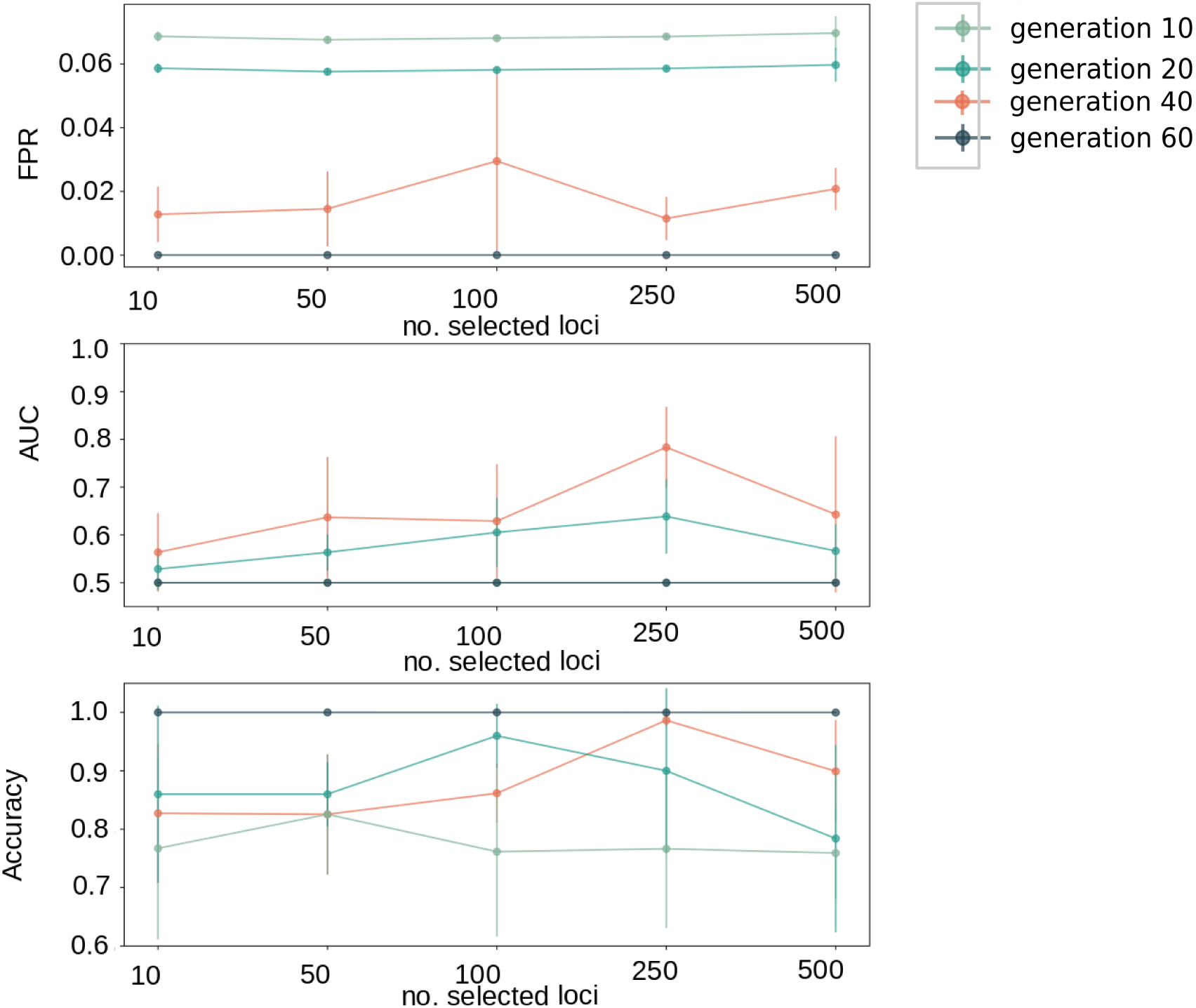
Performance metrics of the combined One Class Support Vector Machines – Fisher’s Exact Test (OCSVM-FET) approach. OCSVM-FET performance metrics at generations 10, 20, 40, and 60 for different numbers of selected loci (n = 10, 50, 100, 250, 500) under strong selection (1.40 Haldane shift). The graph illustrates false positive rate (FPR), area under curve (AUC), and accuracy across the adaptive trajectory. Peak performance occurred at generation 40 with 250 selected loci, demonstrating the lowest FPR, highest AUC (>80%), and near-perfect accuracy (∼99%). This optimal detection window represents the phase where populations have accumulated substantial allele frequency changes but have not yet approached the phenotypic optimum where selection signals become obscured by genetic compensation. The specific timing of this window depends on selection strength and genetic architecture; under different selective regimes, optimal detectability would occur at different generations.

This optimal detection window likely reflects the phase of adaptation where directional selection has driven substantial, coordinated allele frequency changes across many loci (creating a strong collective signal), yet populations have not yet approached the phenotypic optimum where compensatory dynamics and genetic redundancy obscure selection signatures. The specific timing of this window depends on selection strength, initial genetic variation, and trait heritability—under our simulation parameters (strong truncating selection, heritability = 0.8), peak detectability occurred around generation 40. Weaker selection or different genetic architectures would shift this optimal detection window to later or earlier generations, respectively.

This combination of parameters offered the best trade-off between sensitivity and specificity in identifying loci under selection, suggesting OCSVM-FET’s potential as a powerful tool for detecting polygenic adaptation in genomic data, particularly when applied to populations undergoing active directional selection but not yet at evolutionary equilibrium.

### OCSVM-FET Detection Limits Across Selection Regimes

To assess the boundaries of our POC-SVM approach, we evaluated OCSVM-FET performance under medium and weak selection regimes using simulated data. These analyses reveal the method’s detection limits and identify conditions necessary for reliable polygenic adaptation inference. Performance was assessed across evolutionary timescales (10-60 generations) to characterize sensitivity thresholds and improvement trajectories. The performance evaluation under reduced selection pressure revealed the limitations of the OCSVM-FET approach, with both medium and weak selection scenarios yielding modest AUC values (0.55-0.65) and accuracy scores below optimal thresholds (Figure 8). However, clear improvement trends emerged with increasing numbers of selected loci and extended evolutionary timescales

**Fig 8.** OCSVM-FET perfomance limitations and improvement trends across selection regimes. Perfomance metrics for medium (green) and weak (red) selection scenarios are shown across generations. Area under Curve (AUC) values (a), False Positive Rate (FPR) (b) and overall Accuracy scores (c) are displayed. Error bars represents standard deviation across replicates. Temporal effects demonstrate consistent improvement trends, though weak selection remains challenging for reliable detection across most conditions tested.

## Discussion

Our study introduced and validated novel approaches to investigate polygenic adaptation by combining standard statistical methods with machine learning techniques. Our results demonstrate that integrating classic statistics with information-theory-based methods significantly improves detection of polygenic adaptation signals.

### Comparative Performance and Temporal Dynamics of Adaptation

The combined OCSVM-FET method consistently outperformed Fisher’s Exact Test across all tested scenarios. Performance peaked at 250 selected loci, suggesting an optimal balance for our detection approach: beyond 250 loci, individual locus contributions to the phenotype become increasingly marginal, resulting in subtle allele frequency changes that fall below our detection threshold despite ongoing selection. With fewer loci, the molecular signature becomes more pronounced per locus but lacks the characteristic multi-locus pattern that OCSVM leverages for detection. Our simulations revealed that OCSVM-FET’s detection power was particularly strong for substantially polygenic architectures (100-250 loci). At generation 40, populations with 100 and 250 selected loci showed pronounced phenotypic shifts (Figure 5), reflecting substantial adaptive progress toward the phenotypic optimum. Critically, at this evolutionary stage, the molecular signature of selection remained robust: allele frequency changes at selected loci were substantial enough to create a detectable collective pattern, yet individual loci had not yet begun to exhibit the frequency fluctuations characteristic of populations near their adaptive optimum. This “sweet spot” enabled optimal discrimination between selected loci and neutral genomic background. In contrast, oligogenic architectures (10-50 loci) presented a different challenge. While these scenarios showed rapid phenotypic adaptation and strong selection on individual loci, the limited number of selected positions reduced the power of pattern-based detection methods like OCSVM, which rely on identifying collective genomic signatures across many loci. Although individual loci experienced strong selection, the molecular pattern lacked the distributed, multi-locus complexity that OCSVM is designed to capture, resulting in suboptimal detection despite clear phenotypic adaptation.

Conversely, highly polygenic architectures (500 loci) exhibited weak per-locus signals despite substantial phenotypic progress. With selection distributed across so many loci, individual allele frequency changes remained small (typically Δp < 0.15), creating a diffuse molecular signature difficult to distinguish from neutral variation. By generation 40, these populations had achieved only modest phenotypic shifts (Figure 5), reflecting the slower adaptive dynamics of highly polygenic traits, and the cumulative molecular signal—though present—remained too subtle for reliable detection at individual loci or in aggregate patterns.

The temporal dynamics of detection power further illustrate the relationship between phenotypic adaptation and molecular detectability. Peak performance at generation 40 (rather than generation 60) indicates that optimal detection occurs not when populations achieve maximal phenotypic adaptation, but rather during the intermediate phase of directional selection when allele frequency changes are substantial yet consistent. Importantly, phenotypic adaptation and molecular detection signal are not equivalent: a population can show strong phenotypic change through the slow rise of a single large-effect allele (producing weak polygenic detection signal) or through coordinated small shifts at many loci (producing strong polygenic detection signal). Our results demonstrate that OCSVM-FET specifically captures the latter—the molecular signature of polygenic adaptation—rather than phenotypic change per se.

### Trade-offs between methods

Our analysis revealed significant differences among the methods tested. Fisher’s Exact Test performed consistently worst, showing the highest FPR, lowest AUC, and generally lowest accuracy. Combining machine learning approaches with FET enhanced results, with OCSVM-FET demonstrating the best overall performance through consistently low FPR with high accuracy and AUC.

Both OCSVM and NBC were parameter-dependent but offered different practical advantages: OCSVM provided superior detection accuracy at the cost of increased computational demands, while NBC offered faster processing suitable for preliminary analyses or large-scale screening. For studies prioritizing detection power and possessing adequate computational infrastructure, OCSVM-FET is recommended. NBC-FET represents a computationally efficient alternative when resources are limited or rapid exploratory analysis is desired. Our use of real-life *C. riparius* data for parameter optimization [59] ensured calibration to biologically realistic selection patterns rather than idealized simulation scenarios [60], balancing sensitivity to genuine signals with robustness against empirical data noise.

### Genetic Linkage and Hitchhiking Effects

Genetic hitchhiking—where neutral alleles linked to beneficial variants increase in frequency due to physical proximity—can lead to false positive signals in selection detection methods. A critical advantage of our simulation-based validation is that we know the exact positions of truly selected loci, enabling us to distinguish between linked false positives (neutral loci detected due to hitchhiking with nearby selected loci) and unlinked false positives (neutral loci erroneously detected far from any selection target).

Examination of the genomic distribution of false positives revealed that OCSVM-FET predominantly identifies genomic regions genuinely involved in adaptation. The majority of false positive detections occurred in genomic regions harboring selected loci, consistent with hitchhiking effects rather than random misclassification across the genome. This indicates that the method successfully narrows the search space to adaptive regions, even when occasionally flagging linked neutral variants rather than the causal selected loci themselves. From a practical perspective, this performance characteristic is advantageous rather than problematic. In real-world applications—experimental evolution studies or natural population genomics—the initial objective is typically to identify genomic regions under selection, not necessarily to pinpoint individual causal variants Subsequent fine-mapping approaches [61] can then be employed to identify causal variants within the regions flagged by OCSVM-FET. Detection of hitchhiking variants effectively achieves this goal, providing a robust foundation for subsequent fine-mapping through linkage disequilibrium analysis, functional annotation, and experimental validation. Moreover, the distinction between “causal” and “hitchhiking” loci can be biologically ambiguous in polygenic adaptation, where linked variants may contribute to phenotypic variation through epistatic interactions or regulatory effects [62].

### Generalizability

The superior performance of our combined OCSVM-FET approach stems from complementary detection capabilities: OCSVM identifies non-linear patterns in allele frequency change distributions across multiple loci, while FET provides statistical rigor for individual locus significance testing. This combination effectively captures complex polygenic signals while filtering false positives, particularly during intermediate phases of adaptation where directional selection has driven substantial allele frequency changes but populations have not yet reached their phenotypic optimum.

A key advantage of the OCSVM-FET framework is that it operates on single time-point comparisons without requiring replicate populations. While more sophisticated tests such as the Cochran-Mantel-Haenszel (CMH) test can leverage replicate populations to improve power and account for stochastic variation, they fundamentally require such replicates—a constraint that limits their applicability in many experimental and natural contexts. In principle, our simulation replicates could be aggregated and analyzed using CMH instead of FET, potentially improving detection power by explicitly modeling inter-replicate variation. However, we designed our validation to reflect scenarios where single population comparisons are necessary, such as experimental evolution studies with limited replication or natural populations sampled at different time points. Recent methodological advances have further improved population genetic tests by accounting for overdispersion [35], but our OCSVM-based pattern recognition approach complements rather than replaces these advances by detecting collective genomic signatures that may be missed by locus-by-locus statistical tests.

Critical limitations emerge when considering the method’s requirements and assumptions. First, OCSVM-FET requires temporal replicates—genome-wide data from populations sampled at minimum two, better multiple time points during adaptation. While routine in experimental evolution, obtaining such time-series data from natural populations is challenging and often impractical, limiting the method’s application primarily to experimental systems or well-sampled natural populations with documented environmental shifts. Second, our validation was conducted under a specific selection regime (truncating selection with a 1.40 Haldane shift, requiring approximately 40 generations to approach the phenotypic optimum). This selection intensity, while substantial, represents a moderate rather than extreme selective pressure—populations experiencing stronger selection would reach their optimum more rapidly, altering the optimal detection window, whereas weaker selection would require extended evolutionary time for detectable signals to accumulate.

Performance under reduced selection pressure revealed important constraints. The method showed diminished accuracy in weak selection scenarios, where per-locus allele frequency changes remain subtle even after extended evolutionary time, making discrimination from neutral drift increasingly difficult. This fundamental signal-to-noise challenge affects all selection detection methods but is particularly relevant for highly polygenic traits where selection is distributed across many loci. These results suggest that OCSVM-FET is best suited for detecting adaptation under moderate-to-strong selection acting on substantially polygenic traits (100-250 loci), particularly in experimental evolution contexts where temporal sampling is feasible and selection pressures are appreciable. The method’s demonstrated effectiveness across various loci numbers and evolutionary time scales indicates adaptability to diverse genetic architectures and selection intensities within these constraints, providing a valuable tool for the growing field of experimental and real-time evolutionary genomics.

### Future Directions

While our OCSVM-based method represents a significant advance in detecting polygenic adaptation, future studies should validate its performance across diverse genetic architectures and selection intensities. The method could be enhanced through advanced parameter tuning techniques and by incorporating time-series data to improve generalizability across different stages of adaptation. Additionally, developing complementary approaches to characterize the functional impact of identified loci would transform this statistical framework into a more comprehensive tool for understanding the genetic basis of adaptation.

## Declarations

### Ethics approval and consent participate

Not applicable

### Consent for publication

Not applicable

### Availability of data and materials

The dataset used for initial simulation includes unpublished experimental data that are part of pngoing research. Code generatedduring this study is made available on github. Access to unpublished datasets may be granted upon reasonable request and with permission from the data generators.

### Competing interests

The authors declare that they have no competing interests

### Funding

The authors acknowledge funding from the Emergent AI Centre funded by the Carl-Zeiss Stiftung

### Authors’ contributions

C.C. conceived he study, collected and analysed the data, and drafted the manuscript. M.P. and S.G. provided significant intellectual input, supervised the research, and critically revised the manuscript. All authors contributed to the final approval of the manuscript.

## Acknowledgements

We gratefully acknowledge the financial support provided by the Emergent AI Center through the Carl-Zeiss-Stiftung. We extend our sincere thanks to Prof. Dr. Peter Tino, Dr. Pietro Verzelli, and Dr. Daniele Zambone for their valuable insights and guidance. We thank Dr. Hristo Todorov, Dr. Maria Esther Nieto Blázques, Dr. Barbara Feldmeyer and Stephan Weißbach for their careful review of the manuscript. We also acknowledge Dr. Quentin Foucault for conducting the experiments and providing pool-Seq data for parameter tuning.

## Notes

### Competing Interest Statement

The authors have declared no competing interest.

### Summary of Updates

Simulation of weak and medium selection has be added and analysed. The new analysis supports the results of the previous version.

