## Supplement for "Enhancing Detection of Polygenic Adaptation: A Comparative Study of Machine Learning and Statistical Approaches Using Simulated Evolve-and-Resequence Data"

#### Material and Method

#### Simulations

MimicrEE2's quantitative trait (QT) mode was employed to simulate rapid polygenic adaptation. This mode first computes phenotypic values for each individual based on the effect sizes of the SNPs and environmental variance, then performs truncating selection, culling individuals with the most extreme phenotypic values. This approach allows for the simulation of complex adaptive events under various evolutionary scenarios.

Our simulation pipeline started with the generation of a haplotype input file using individual whole genome sequencing data from 66 *C. riparius* individuals, that were conducted during the study of Foucault et. al. [1]. We utilized the non-model organism pipeline for GATK4 [2] (version 4.5.0.0) to create haplotypes from individual sequencing fastq files. These empirical haplotype data served as the ancestral generation for subsequent simulations, providing a biologically relevant starting point specific for *C. riparius*.

Using this haplotype file, we conducted a total of 20 distinct simulation setups, which were designed to explore two key variables:

1. Time under selection: We simulated four different periods under selection - 10, 20, 40, and 60 generations. This range allowed us to observe how adaptation patterns developed over different time scales.
2. Number of loci under selection: We varied the number of loci experiencing selection, testing five different quantities - 10, 50, 100, 250, and 500. This variation helps us understand how the number of genes involved in adaptation affects detection methods, ranging from oligogenic over mildly to highly polygenic adaptation.

For each combination of generation time and number of selected loci, we ran 10 replicates to account for stochastic variations in evolutionary processes. This design resulted in a total of 200 simulation runs (4 generation times × 5 loci numbers × 10 replicates) (Tab 2)

Using *qt* mode, we modelled a scenario of strong selection by specifying a range of effect sizes for the loci under selection. The effect size represents the magnitude of influence a particular allele has on the phenotypic value of an individual. MimicrEE2 uses these effect sizes in the following way: Firstly, for each locus under selection, an effect size is randomly chosen from our specified range (in our case: 1.5 to 3.5). Secondly, the phenotypic value of an individual is calculated as the sum of the products of each locus's effect size and its allele frequency, plus random environmental noise. Lastly, the initial allele frequencies provided together with the effect size are used as the starting point for the simulation, representing the genetic makeup of the population at generation 0. We chose a high effect size in range of 1.5 to 3.5 allowing us to observe significant changes within a relatively small number of generations, which is computationally efficient and helps in detecting selection signals more clearly.

We generated a selected loci file using Python (version 3.8.13) and the library pandas (version 2.0.3) [3], which specifies the loci under selection and incorporates the effect sizes of the loci contributing to a quantitative trait. The script randomly selected 10, 50, 100, 250, and 500 positions with a starting allele frequency provided by the effect-file. In our case, we chose allele frequencies ranging from 0.01 to 0.3.

The recombination map was generated using Python and the library pandas. We allowed the recombination rate to vary between 0.1 and 4 centimorgans, encompassing a range of possible recombination rates that most likely reflect the natural variation observed in *C. riparius* populations [4].

Another key parameter was heritability, which was set to 0.8, indicating a strong genetic component in the trait under selection. A truncating selection model was applied with a factor of 0.5, i.e. removing individuals with phenotypic values below the 50th percentile.

The output was stored in sync-file format to facilitate comparison with real-life experimental data. We utilized the R package poolSeq for subsequent analysis of allele frequencies across generations.

This simulation design allowed us to model rapid adaptation under strong selection pressure. By incorporating real genetic data from *C. riparius* and varying key parameters such as selection coefficient and recombination rate, we aimed to capture a range of evolutionary scenarios, providing a robust test for our analytical methods in identifying polygenic adaptation patterns (parameters: java -jar mim2-v206.jar/ qt/ --haplotypes-g0 haplotype-file.mimhap/ --recombination-rate recombinationrate-file/ --effect-size (--n 10, 50, 100, 250, 500/ --f 0.15/ --effect-file )/ --heritability 0.8/ --snapshots 10,20,40,60 / --replicate-runs 10 / --selection-regime 0.5 / --threads 6 --output-sync output.sync.gz /).

**Table 2. Schematic construction of simulated data set by Mimicree2.**

| | | |
|---|---|---|
| Generation 10 | 10 loci | 10 replicates |
| | 50 loci | 10 replicates |
| | 100 loci | 10 replicates |
| | 250 loci | “ |
| | 500 loci | “ |
| Generation 20 | 10 loci | “ |
| | 50 loci | “ |
| | 100 loci | “ |
| | 250 loci | “ |
| | 500 loci | “ |
| Generation 40 | 10 loci | “ |
| | 50 loci | “ |
| | 100 loci | “ |
| | 250 loci | “ |

| | | |
|---|---|---|
| | 500 loci | “ |
| Generation 60 | 10 loci | “ |
| | 50 loci | “ |
| | 100 loci | “ |
| | 250 loci | “ |
| | 500 loci | “ |

For each generation 10, 20, 40 and 60 and each selected loci setting (n= 10, 50, 100, 250, 500) 10 replicate pools were created, originating from 66 individual sequencing files [1].

#### Phenotypic value calculation

While MimicrEE2 does not directly output phenotypic values, we derived these values from the simulation outputs using a custom Python script (version 3.8.13). The phenotypic values were calculated using the allele frequencies from the MimicrEE2 output and the effect sizes provided as input to the algorithm.

The calculation of phenotypic values was performed as follows:

1. Effect sizes were randomly sampled from the input effect size file for each locus under selection, allowing for replacement to match the number of loci in each scenario.

2. The raw phenotypic value for each generation was computed using the formula:
   **Raw Phenotypic Value = Genetic Component + Environmental Component**
   Where:

   - Genetic Component = Σ(p_i × effect_i) for all loci i
   - p_i = allele frequency at locus i (from MimicrEE2 output)
   - effect_i = effect size at locus i (randomly sampled from uniform distribution between 1.5 and 3.5)
   - Environmental Component = random noise drawn from a normal distribution with variance scaled to maintain heritability of 0.8

3. For each scenario, a phenotypic optimum was calculated as the sum of all positive effect sizes (Optimum = Σ(max(effect_i, 0))), representing the theoretical maximum phenotypic value achievable when all beneficial alleles reach fixation.

4. To enable comparison of adaptive trajectories across scenarios with different numbers of selected loci, raw phenotypic values were normalized relative to each scenario's evolutionary trajectory: Normalized Phenotypic Value = (Raw Value - Ancestral Mean) / (Optimum - Ancestral Mean) This normalization scales values such that 0 represents the ancestral population mean and 1.0 represents achievement of the scenario-specific phenotypic optimum (complete fixation of all beneficial alleles for that particular genetic architecture). Values between 0 and 1

indicate partial progress toward the optimum. While absolute phenotypic values differ across scenarios due to varying numbers of loci and their associated effect sizes, this normalization allows comparison of the relative rate and extent of adaptive progress toward each scenario's evolutionary optimum.

5. This process was repeated for each scenario (10, 50, 100, 250, and 500 loci) using the corresponding allele frequency files generated by MimicrEE2. For visualization purposes, we present the mean normalized phenotypic values across generations at key timepoints (generations 10, 20, 40, and 60).

#### Bayes' Theorem

The derivation of Bayes' theorem forms the basis for the following workflow and formulas for the Naive Bayesian Classifier, that were described in [5] [6] [7]:

The aim is to model three classes, one class for the non-anomalous data, one class for the anomalous data increasing in AF over time and another class for anomalous data with decreasing AF over time.

We assume there are k classes, $C_1, C_2, ..., C_k$. Each sample is represented by an n-dimensional vector, $X = \{x_1, x_2, ..., x_n\}$, signifying n measured values of n attributes $A_1, A_2, ..., A_n$, respectively. Given a sample X, the classifier predicts that X belongs to the class with the highest probability conditioned on X. In other words, X is predicted to belong to class $C_i$ if and only if:

$$P(C_i \vee X) > P(C_j \vee X) \text{ for } 1 \leq j \leq m, \text{ where } j \neq i \quad (1)$$

Thus, we identify the class that maximizes $P(C_i|X)$. The class $C_i$ for which $P(C_i|X)$ is maximized is called the maximum a posteriori hypothesis. According to Bayes' theorem:

$$P(C_i \vee X) = [P(X \vee C_i) \times P(C_i)] / P(X) \quad (2)$$

Evaluating $P(C_i|X)$ for data sets with numerous attributes can be computationally resource-intensive. To compensate for this, the naive assumption of class conditional independence is employed. This assumption implies that the attribute values are conditionally independent of one another given the class label of the sample. Mathematically, this translates to:

$$P(X \vee C_i) \approx \prod_{k=1}^{n} P(x_k \vee C_i) \quad (3)$$

When attribute $A_k$ is continuous-valued, it is common to assume that the values follow a Gaussian distribution with a mean μ and a standard deviation σ, defined as:

$$g(x, mu, sigma) = \frac{1}{\sqrt{2 pi \cdot sigma}} \times \exp\left(\frac{-(x - mu)^2}{2 sigma^2}\right) \quad (4)$$

in one-dimensional space. In 2D the mean μ becomes an n-dimensional vector and the deviation σ becomes an n × n covariance matrix Σ:

$$g(x; mu, Sigma) = \frac{1}{(2\pi)^{n/2} \cdot |Sigma^{1/2}|} \cdot \exp\left(-1/2\,(x-mu)^{T}\, Sigma^{-1}\,(x-mu)\right) \quad (5)$$

To predict the class label of X, P(X|$C_i$) × P($C_i$) is evaluated for each class $C_i$. The classifier predicts that the class label of X is $C_i$ if and only if it maximizes P(X|$C_i$) × P($C_i$).

Different settings for μ and Σ will vary the modelling of the data, as displayed with toy data by Figure S1.

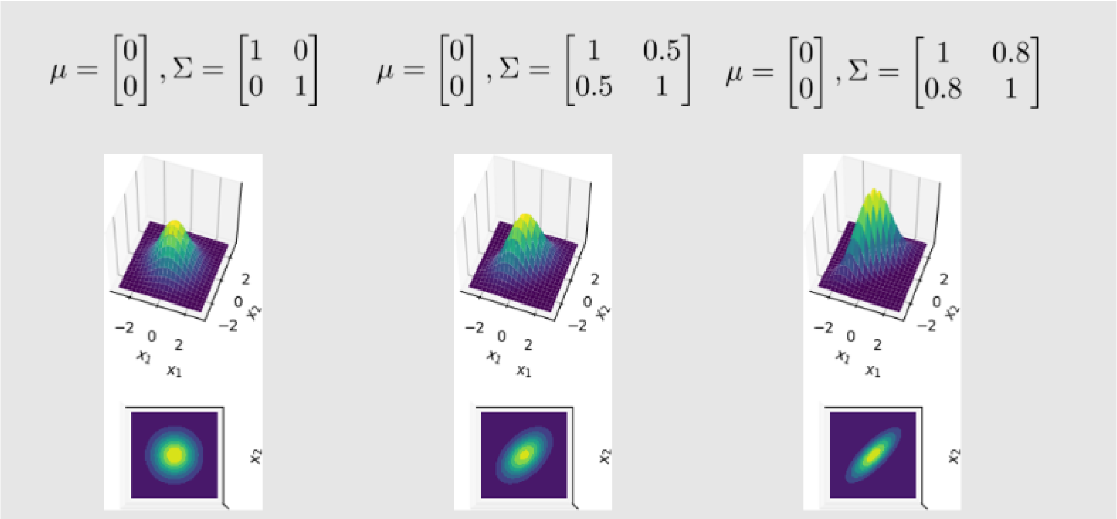


**Fig S1. Example illustration of the effects of different parameter settings for the covariance matrix on the multivariate probability density function.** The modelling of the data changes depending on the values of the covariance parameters.

#### Parameter Fine Tuning

The key parameters μ and Σ for NBC, as well as ν and γ for OCSVM were optimized using real-life and previously published datasets by Pfenninger et al. [8]. This data, originating from an experiment on *C. riparius*, captured genome-wide allele frequency changes before and after a cold snap event, suggesting a potential rapid, polygenic adaptation in the population. DNA extraction, sequencing, trimming, mapping, filtering of reads and SNP calling was performed as described in [8]. To infer long-term selection regimes, Tajima's D was calculated. The temporal behavior of candidate loci was compared to randomly selected SNPs to assess their correlation. In this study, several genetic markers were identified with significant allele frequency shifts before and after the cold snap, indicating potential selection events. These loci were used as true positives to fine-tune the parameters for the introduced methods NBC and OCSVM. The optimized parameter settings were then evaluated for accuracy, false positive rate (FPR) and area under curve (AUC) using simulated data.

While the original study identified 19 SNPs as candidates for selection, these were consolidated into 10 independent loci due to linkage on the same scaffolds [8]. For our parameter fitting process, we used these 10 independent loci as true positives, as they represent distinct genomic regions potentially affected by selection and are likely physically unlinked.

To optimize OCSVM parameters, we performed a grid search over ν values ranging from 0.001 to 1 and γ values from 0.01 to 1. For NBC, we explored various combinations of μ and Σ values, ensuring the resulting covariance matrices remained valid positive semi-definite. The settings for these parameters can be found in the Supplementary Fig S2. The grid search aimed to maximize performance across accuracy, FPR and AUC. These metrics were chosen to balance the algorithm's ability to correctly identify true positives while minimizing false detections, crucial for detecting subtle signals of polygenic adaptation.

The optimized parameters were then applied to simulated data mimicking various evolutionary scenarios, including different generations (10, 20, 40, 60) and numbers of loci under selection (10, 50, 100, 250, 500). This allowed us to assess how well the parameters, tuned on real data, performed across a range of potential adaptive scenarios.

Using the 10 independent loci as true positives for parameter tuning aligns with the expectation that polygenic adaptation often involves multiple, unlinked genomic regions. This approach allowed us to capture the diversity of allele frequency changes associated with complex adaptive processes.

### Results

**Tab S1. Parameter optimization for OCSVM and NBC algorithms.**

a

| Setting | $\nu$ | $\gamma$ | FPR | AUC | ACCURACY |
|---|---|---|---|---|---|
| 1 | 0.001 | 0.01 | 0.001 | 0.5 | 0.999 |
| 2 | 0.001 | 0.05 | 0.001 | 0.499 | 0.999 |
| 3 | 0.001 | 0.5 | 0.002 | 0.541 | 0.998 |
| 4 | 0.001 | 1 | 0.003 | 0.624 | 0.997 |
| 5 | 0.005 | 0.01 | 0.005 | 0.914 | 0.995 |
| 6 | 0.005 | 0.05 | 0.005 | 0.872 | 0.995 |
| 7 | 0.005 | 0.5 | 0.006 | 0.914 | 0.994 |
| 8 | 0.005 | 1 | 0.006 | 0.914 | 0.994 |
| 9 | 0.01 | 0.01 | 0.010 | 0.912 | 0.990 |
| 10 | 0.01 | 0.05 | 0.010 | 0.995 | 0.990 |
| 11 | 0.01 | 0.5 | 0.011 | 0.995 | 0.989 |
| 12 | 0.01 | 1 | 0.011 | 0.911 | 0.989 |
| 13 | 0.013 | 0.01 | 0.013 | 0.910 | 0.987 |
| 14 | 0.013 | 0.05 | 0.013 | 0.993 | 0.987 |
| 15 | 0.013 | 0.5 | 0.014 | 0.993 | 0.986 |
| 16 | 0.013 | 1 | 0.012 | 0.910 | 0.988 |
| 17 | 0.02 | 0.01 | 0.020 | 0.990 | 0.980 |
| 18 | 0.02 | 0.05 | 0.020 | 0.990 | 0.980 |
| 19 | 0.02 | 0.5 | 0.019 | 0.991 | 0.981 |

| | | | | | |
|---|---|---|---|---|---|
| 20 | 0.02 | 1 | 0.020 | 0.990 | 0.980 |
| 21 | 0.1 | 0.01 | 0.1 | 0.950 | 0.900 |
| 22 | 0.1 | 0.05 | 0.101 | 0.950 | 0.899 |
| 23 | 0.1 | 0.5 | 0.1 | 0.950 | 0.900 |
| 24 | 0.1 | 1 | 0.1 | 0.950 | 0.900 |

b

| Seeting | $\mu_1$ | $\Sigma_1$ | $\mu_2/\mu_3$ | $\Sigma_2/\Sigma_3$ | FPR | AUC | AUCCURAY |
|---|---|---|---|---|---|---|---|
| 1 | 0.0, 0.0 | 0.15, 0.1 | 0.1, 0.9 | 0.25, 0.2 | 0.004 | 0.998 | 0.996 |
| 2 | 0.0, 0.0 | 0.25, 0.18 | 0.2, 0.7 | 0.2, 0.199 | 0.003 | 0.874 | 0.997 |
| 3 | 0.0, 0.0 | 0.15, 0.1 | 0.3, 0.4 | 0.35, 0.25 | 0.115 | 0.943 | 0.885 |
| 4 | 0.0, 0.0 | 0.2, 0.1 | 0.3, 0.5 | 0.4, 0.2 | 0.073 | 0.922 | 0.927 |
| 5 | 0.0, 0.0 | 0.3, 0.1 | 0.3, 0.6 | 0.25, 0.2 | 0.320 | 0.840 | 0.680 |
| 6 | 0.0, 0.0 | 0.25, 0.2 | 0.3, 0.8 | 0.15, 0.1 | 0.027 | 0.986 | 0.973 |
| 7 | 0.0, 0.0 | 0.2, 0.15 | 0.35, 0.6 | 0.3, 0.2 | 0.065 | 0.967 | 0.935 |
| 8 | 0.0, 0.0 | 0.3, 0.2 | 0.5, 0.4 | 0.35, 0.3 | 0.374 | 0.813 | 0.626 |
| 9 | 0.0, 0.0 | 0.25, 0.2 | 0.6, 0.3 | 0.2, 0.15 | 0.087 | 0.957 | 0.913 |
| 10 | 0.0, 0.0 | 0.15, 0.1 | 0.7, 0.2 | 0.25, 0.2 | 0.020 | 0.990 | 0.980 |

(a) Tested parameter settings (v and γ) for the OCSVM algorithm with their respective performance metrics. (b) Tested parameter settings for NBC, showing mu and sigma values for three approximations along with their performance metrics

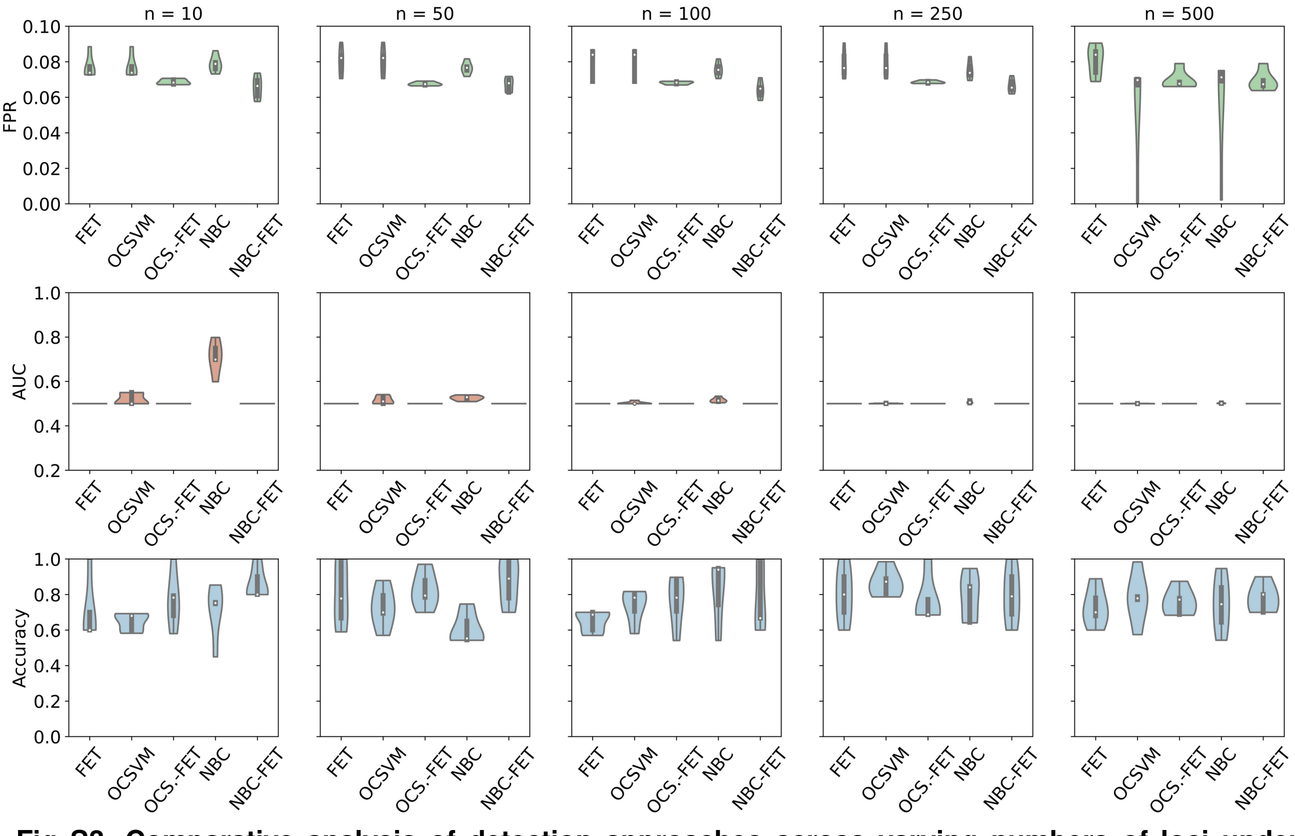


**Fig S3. Comparative analysis of detection approaches across varying numbers of loci under selection at generation 10.** The graph displays False Positive Rate (FPR) (first column graphs, green), Area Under the Curve/Receiver Operating Characteristic (AUC/ROC) (second column graphs, red), and

Accuracy (third column graphs, blue for FET, OCSVM, OCSVM-FET, NBC, and NBC-FET. Each row represents the number of loci under selection (n = 10, 50, 100, 250, 500).

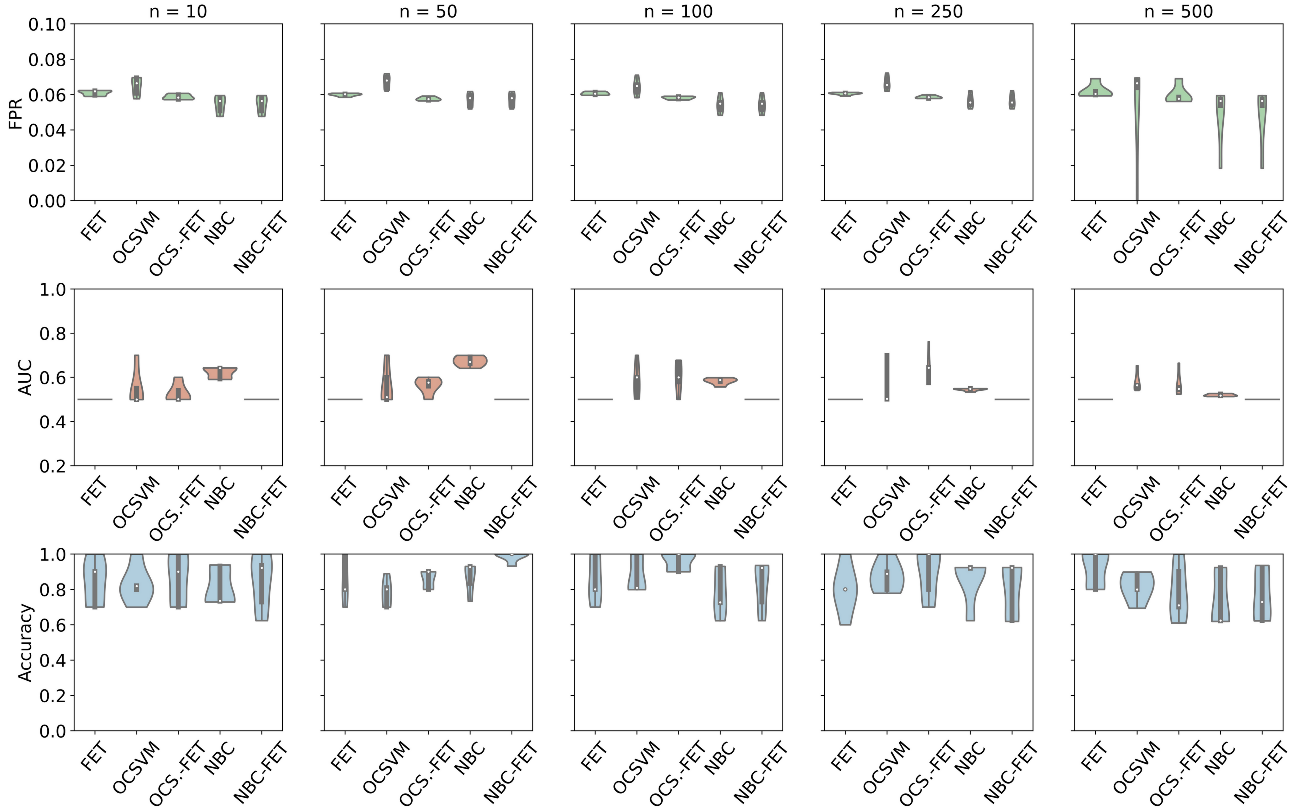


**Fig S4. Comparative analysis of detection approaches across varying numbers of loci under selection at generation 20.** The graph displays False Positive Rate (FPR) (first column graphs, green), Area Under the Curve/Receiver Operating Characteristic (AUC/ROC) (second column graphs, red), and Accuracy (third column graphs, blue for FET, OCSVM, OCSVM-FET (OCS.-FET), NBC, and NBC-FET. Each row represents the number of loci under selection (n = 10, 50, 100, 250, 500).

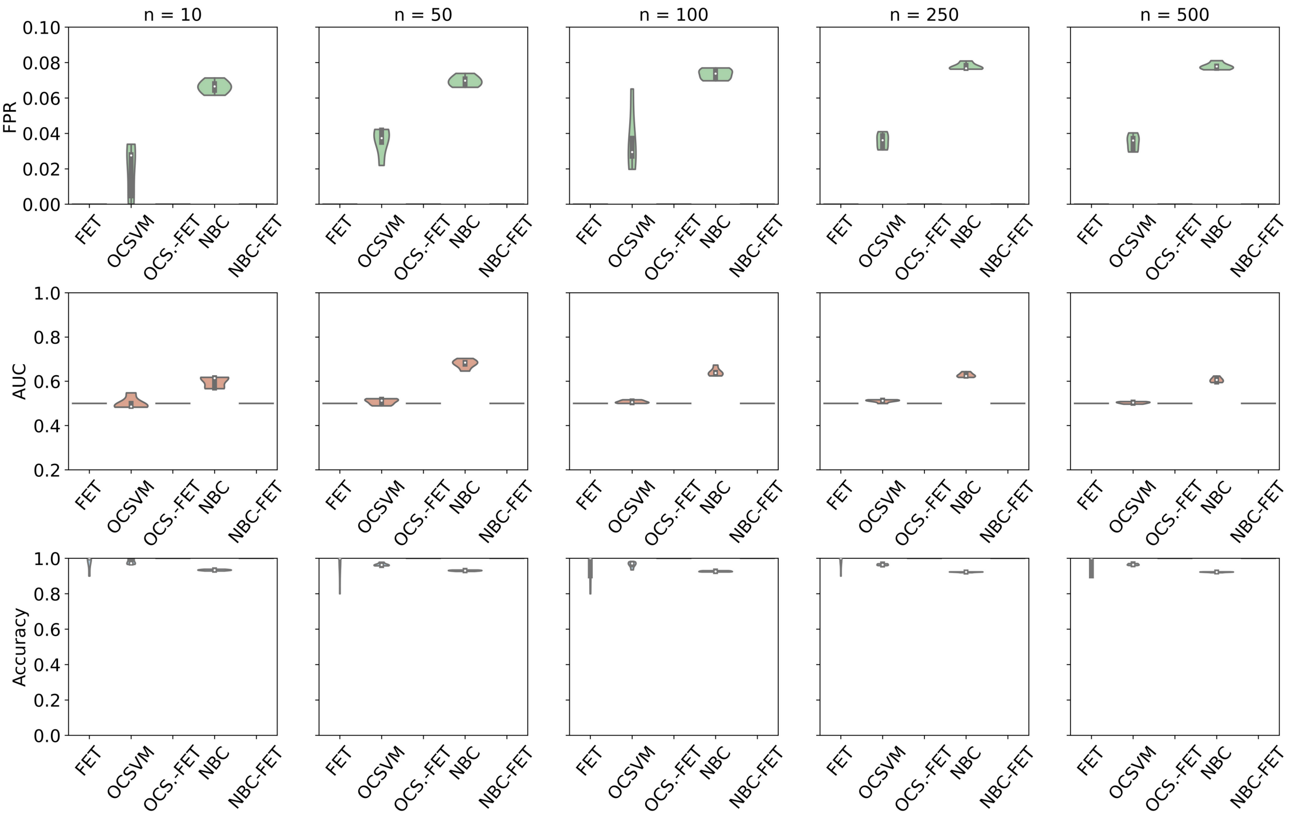


**Fig S5: Comparative analysis of detection approaches across varying numbers of loci under selection at generation 60.** The graph displays False Positive Rate (FPR) (first column graphs, green), Area Under the Curve/Receiver Operating Characteristic (AUC/ROC) (second column graphs, red), and Accuracy (third column graphs, blue for FET, OCSVM, OCSVM-FET (OCS.-FET), NBC, and NBC-FET. Each row represents the number of loci under selection (n = 10, 50, 100, 250, 500).

[5] I. Wickramasinghe and H. Kalutarage, “Naive Bayes: applications, variations and vulnerabilities: a review of literature with code snippets for implementation.,” *Soft*

*Comput.*, vol. 25, no. 3, pp. 2277–2293, Sep. 2020, doi: 10.1007/S00500-020-05297-6.

[6] K. Kaushik *et al.*, “Multinomial Naive Bayesian Classifier Framework for Systematic Analysis of Smart IoT Devices,” *Sensors (Basel).*, vol. 22, no. 19, p. 7318, Oct. 2022, doi: 10.3390/S22197318.

[7] R. R. Yager, “An extension of the naive Bayesian classifier,” *Inf. Sci. (Ny).*, vol. 176, no. 5, pp. 577–588, Mar. 2006, doi: 10.1016/J.INS.2004.12.006.

[8] M. Pfenninger, Q. Foucault, A. M. Waldvogel, and B. Feldmeyer, “Selective effects of a short transient environmental fluctuation on a natural population,” *Mol. Ecol.*, vol. 32, no. 2, pp. 335–349, Jan. 2023, doi: 10.1111/MEC.16748.
